# Ocular hypertension drives remodeling of AMPA receptors in select populations of retinal ganglion cells

**DOI:** 10.1101/743559

**Authors:** Asia L. Sladek, Scott Nawy

**Affiliations:** Truhlsen Eye Institute, Department of Ophthalmology and Visual Sciences, University of Nebraska Medical Center, Omaha, NE 68198

## Abstract

AMPA receptors in the CNS are normally impermeable to Ca^2+^ but aberrant expression of Ca^2+^-permeable AMPA receptors (CP-AMPARs) occurs in pathological conditions such as ischemia or epilepsy, or in degenerative diseases such as ALS. Here we show that select populations of retinal ganglion cells (RGCs) similarly express high levels of CP-AMPARs in a mouse model of glaucoma. CP-AMPAR expression increased dramatically in both α On and α transient Off RGCs, and this increase was prevented by genomic editing of the GluA2 Q/R site. α On RGCs with elevated CP-AMPAR levels displayed profound synaptic depression, which was reduced by selectively blocking CP-AMPARs, buffering Ca^2^+ with BAPTA, or with the CB1 antagonist AM251, suggesting that depression was mediated by a retrograde transmitter which might be triggered by influx of Ca^2^+ through CP-AMPARs. Thus OHT alters the composition of AMPARs and modulates patterns of synaptic activity in select populations of RGCs.

## Introduction

Glaucoma is a neurodegenerative disease of retinal ganglion cells (RGCs) often associated with elevated intraocular pressure (IOP). Several mouse models have been developed to induce IOP, including injection of microspheres which reduce aqueous outflow through the trabecular meshwork, causing a long lasting increase in ocular pressure (Sappington et al., 2010). Within one week following elevation of IOP, there are a number of morphological and functional changes in RGCs including axonal transport (Buckingham et al., 2008; Crish et al., 2010; Calkins, 2012; Ward et al., 2014), dendritic morphology (Della Santina et al., 2013; El-Danaf and Huberman, 2015), decreases in synaptic responses, the patterns of light-driven input to RGCs (Holcombe et al., 2008; Frankfort et al., 2013; Chen et al., 2015; El-Danaf and Huberman, 2015; Pang et al., 2015) and membrane excitability (Risner et al., 2018). However, it is currently unclear whether these previously described changes contribute directly to RGC death, or whether other yet to be described changes are critical.

There has been a long standing interest in the contribution of aberrant glutamate receptor expression to the underlying etiology of glaucoma (reviewed in Almasieh et al., 2012). While early studies focused on the contribution of highly Ca^2+^-permeable NMDA receptors, recent studies have focused on a for a role of Ca^2+^-permeable AMPA receptors (CP-AMPAR) (Wang et al., 2014; Cueva Vargas et al., 2015; Wen et al., 2018). CP-AMPARs are upregulated at synapses in response to a number of pathological conditions such as ischemic insult (Liu et al., 2004; Noh et al., 2005; Kwak and Weiss, 2006), ALS (Kwak et al., 2010; Yamashita et al., 2013), Alzheimer’s and Parkinson’s disease (Kobylecki et al., 2010; Whitehead et al., 2017), and drug addiction (Shukla et al., 2017). Permeability to Ca^2+^ is determined by the absence or presence of the GluA2 subunit. This subunit contributes a positively charged arginine residue to the channel pore, preventing passage of Ca^2+^ and Zn^2+^ (Cull-Candy et al., 2006; Bowie, 2012). The presence of this arginine depends on the RNA editing enzyme ADAR2 (Lomeli et al., 1994; Higuchi et al., 2000; Horsch et al., 2011). In the unedited form of GluA2, a neutral glutamine is expressed, allowing for Ca^2+^ permeation. Thus AMPARs are permeable to Ca^2+^ if they lack GluA2, or contain an unedited form of the subunit. In response to pathological conditions, it appears that AMPA receptors can be remodeled to increase Ca^2+^ permeability via either mechanism.

Under normal conditions, CP-AMPAR also play important roles in the induction and maintenance of synaptic plasticity in a number of brain regions. In cerebellum, where it was first described, high frequency stimulation of presynaptic parallel fibers drives the rapid replacement of CP-AMPARs with Ca^2+^-impermeable AMPARs (CI-AMPAR). The initial event that triggers this plasticity is Ca^2+^ influx through the CP-AMPAR itself (Liu and Cull-Candy, 2000). Insertion of CP-AMPARs is a critical step in consolidation of fear driven memories (Clem and Huganir, 2010; Liu et al., 2010; Rao-Ruiz et al., 2011; Hong et al., 2013). One consequence of a switch from CI-to CP-AMPAR is a change in postsynaptic excitability (Savtchouk and Liu, 2011; Liu and Savtchouk, 2012), but local increases in Ca^2+^ via influx through CP-AMPARs may have other consequences as well.

Here we show that two weeks of ocular hypertension (OHT) is sufficient to remodel AMPARs in α On and α transient Off, but not α sustained Off RGCs. Amongst the alpha type RGCs, the Off transient type appears more vulnerable than the others, although there is some subtype variability depending upon the parameters that are being measured (Della Santina and Ou, 2017). AMPARs displayed increased voltage-dependent block by spermine, consistent with increased CP-AMPARs expression (Bowie and Mayer, 1995; Donevan and Rogawski, 1995; Kamboj et al., 1995; Koh et al., 1995). The presumed changes in AMPAR calcium permeability induced by OHT were not observed in a mouse line in which the GluA2 editing is built-in using transgenic substitution of arginine for glutamine at the Q/R site, suggesting that the reorganization is accomplished by reduced RNA editing of GluA2, rather than removal of the subunit.

We also find, using an optogenetic approach, that OHT decreases synaptic gain at bipolar to α On RGC synapse. Interesting, a decrease in synaptic gain was observed previously in a variant of CP-AMPAR plasticity in which Ca^2+^ influx through NMDA receptors drives replacement of CI-AMPARs with CP-AMPARs in the same type of RGC (Jones et al., 2012). Thus two different experimental conditions, chronic elevation of ocular pressure, or acute NMDA receptor activation, converge onto the same cell type to elevate CP-AMPARs, and decrease synaptic gain. We explore the possibility that CP-AMPAR expression and synaptic gain are linked, with CP-AMPARs providing a route of Ca^2+^ influx to activate a retrograde messenger that reduces transmitter release from the presynaptic bipolar cell.

## Results

### α transient and sustained Off RGCs express predominantly Ca^2+^-impermeant AMPARs in normal retina

We first targeted α Off RGCs, identified by their large size, slightly oval shape, and characteristic light responses, as described below. Two classes of α Off RGCs, the α Off transient RGCs and the α Off sustained could be distinguished using several criteria. The α Off transient RGC expressed a robust T type Ca^2+^ current (Fig. 1A, left), as described previously (Margolis and Detwiler, 2007; van Wyk et al., 2009; Murphy and Rieke, 2011). The amplitude distribution of T type Ca^2+^ current that was measured in α RGCs could be well fit with a single Gaussian, suggesting that the distribution describes a single population of cells (Fig. 1D). α Off transient RGCs responded to termination of dim light (1 s stimulus of 490 nm, intensity of ~0.7 to 7 Rh*/rod) with a large inward current which rapidly decayed to a small sustained dark current (Fig. 1A, right). Conversely, α Off sustained RGCs lacked T type Ca^2+^ currents (Fig. 1B, left), and they typically had a larger sustained inward current in darkness compared with transient cells, as might be expected if this ongoing excitatory input contributes to spike generation in darkness (Fig. 1B). For α Off sustained RGCs, the sustained dark current (thin to dashed lines) accounted for 29±5% of the total peak to peak current (thin to thick lines), compared to 13±3% in α Off transient RGCs (Fig. 1C). In some cells, identification of cell type was confirmed by filling with Neurobiotin and examining the layer of termination of the dendrites (van Wyk et al., 2009; Bleckert et al., 2014).

**Figure 1.**
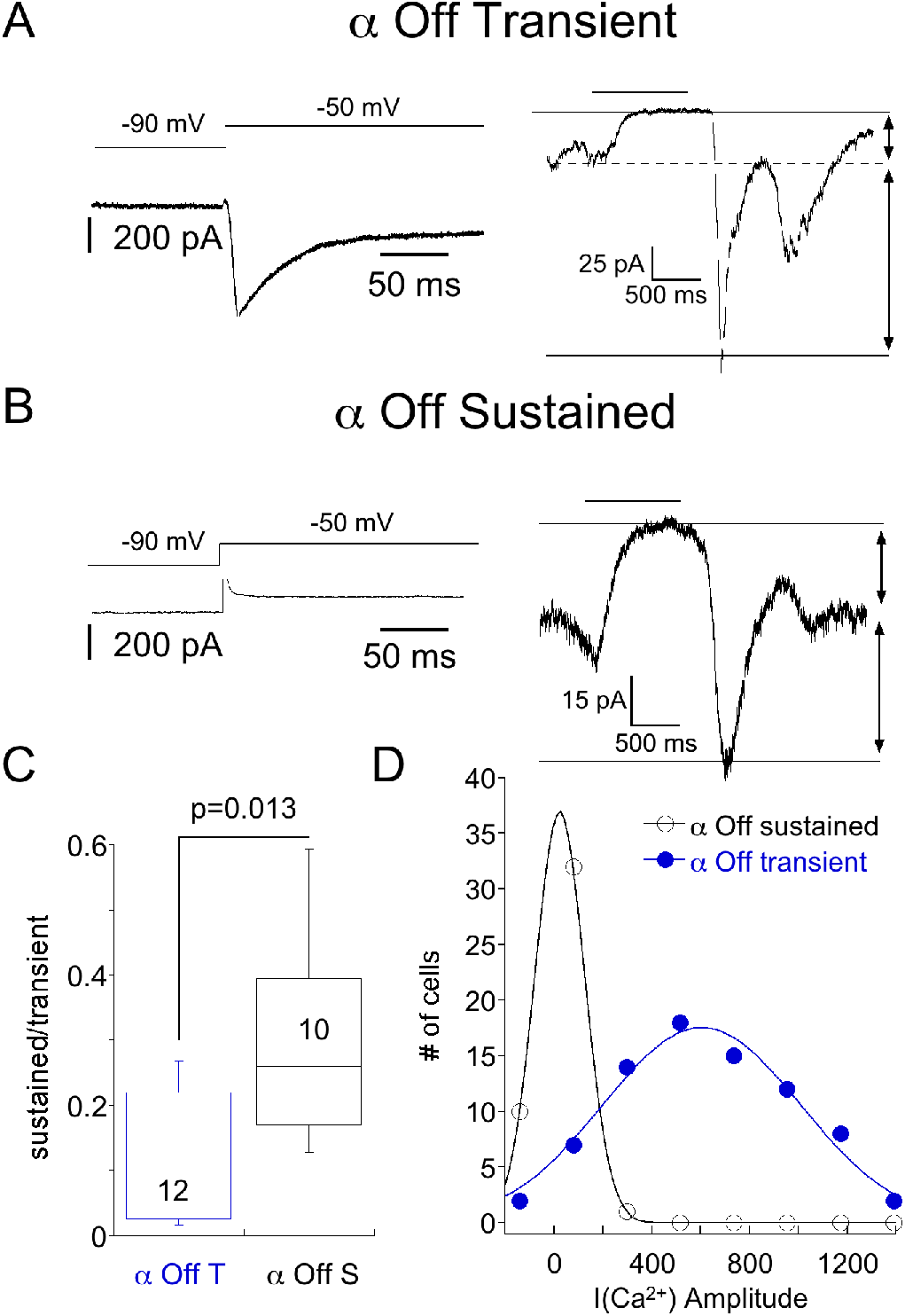
Responses of α Off sustained and transient RGCs to steps of dim light. A. Left, Ca^2+^ current evoked by voltage step from −90 to −50 mV. Right, response of the same cell to a 490 nm light (6.8 R*/rod). The light-suppressed continuous current is indicated by the thin and dashed lines, and the transient response at light off is bounded by the broken and thick solid line. B. Example of an Off cell lacking a T-type Ca^2+^ current. In these this cell type, the transient current was smaller in amplitude compared with α Off transient RGCs. C. Box plot summarizing the ratio of the sustained, light suppressed current to the transient current observed at light off for presumed α Off transient and α Off sustained RGCs. D. Histogram of Ca^2+^ current amplitudes for presumed α Off transient and α Off sustained RGCs. Continuous lines are Gaussian fits with peaks at 604 pA and 23 pA, respectively.

We determined if CP-AMPARs contribute synaptic transmission from bipolar cells to either α Off sustained or transient RGCs by measuring the IV relationship of the light response in both types of cells. NMDA receptors were blocked with D-APV (50 μM), and feed forward inhibition was blocked with strychnine and picrotoxin to avoid contamination of the I-V relationship with non-AMPA currents. Spermine (500 μM), which selectively blocks CP-AMPARs at positive, but not negative holding potentials (Bowie and Mayer, 1995; Kamboj et al., 1995; Koh et al., 1995), was included in the pipet solution. An example of the light response of an α transient Off RGC at 3 holding potentials is shown in figure 2A. AMPA current was measured at the cessation of light (filled circle), when transmitter release from bipolar cells reaches a peak. If CP-AMPARs contribute to the light response, they should be open at negative voltages, but blocked at positive voltages, resulting in the inward rectification of the IV relation. However, this was not observed in RGCs taken from eyes with normal ocular pressure, as the amplitude of the light response was not diminished at positive voltages.

**Figure 2.**
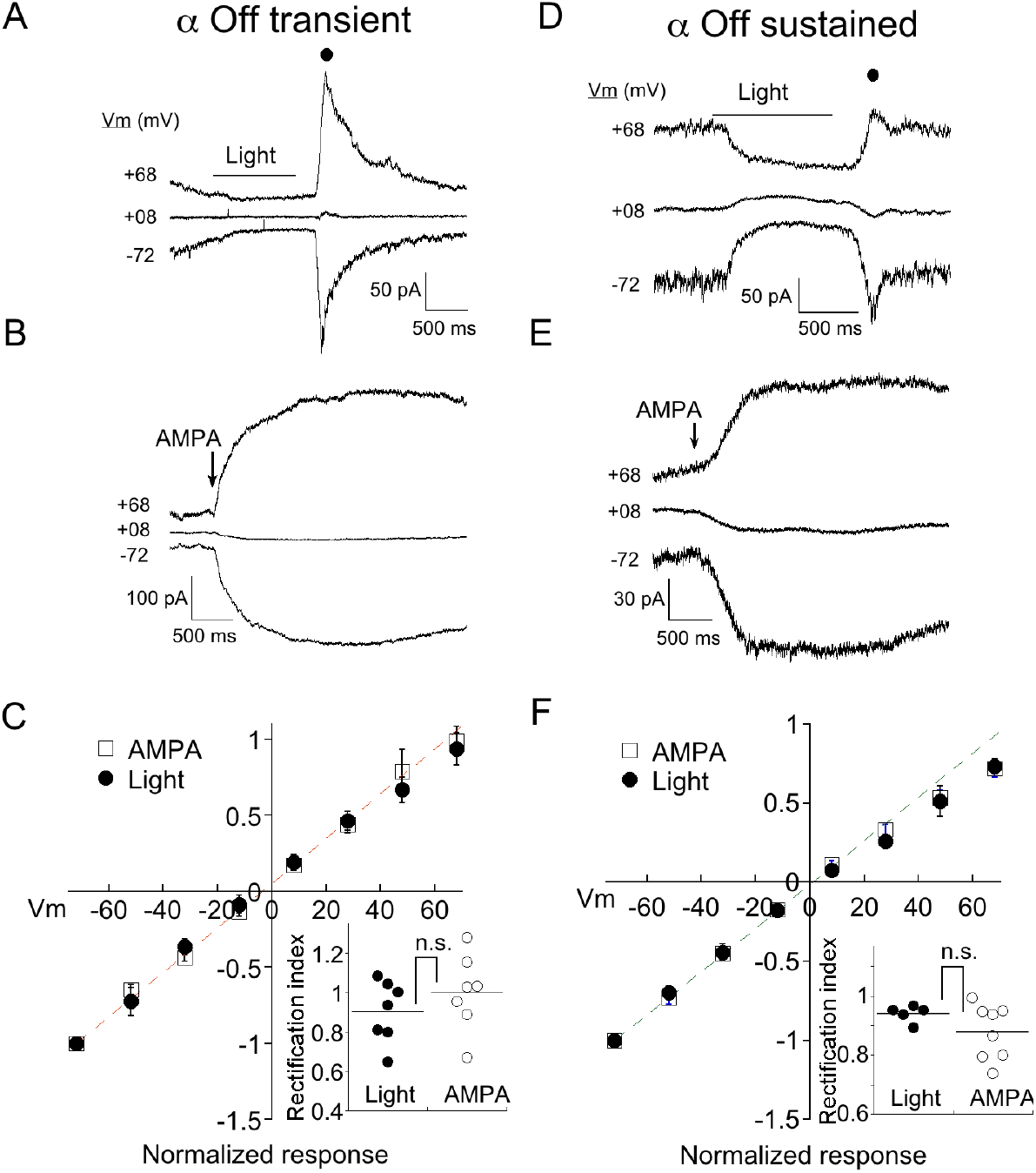
α Off transient and sustained RGCs preferentially express CI-AMPARs under normal conditions. A. Response of an α Off transient RGC to a 1 second 490 nm light delivering 6.8 R*/rod at three holding potentials. Spermine (500 μM) is included in the pipet solution here and in all subsequent figures. B. Response of a different α Off transient cell to a 100 ms application of 100 μM AMPA. C. Averaged IV relation of AMPA currents evoked with either light or exogenous AMPA. Dashed line indicates the linear fit to the current at negative holding potentials. Inset: The rectification index, calculated from the IV relation for each cell (see methods). The lack of significant rectification in the presence of intracellular spermine, probed with either light or AMPA, suggests that few CP-AMPARs are expressed in eyes with normal ocular pressure. D-F. Exemplar responses of α Off sustained RGCs to light and AMPA, and the normalized mean IV relations. As with α Off transient RGCs, the IV relations of AMPA currents in α Off sustained RGCs were linear regardless of the method used to activate channels.

As a second method for estimating CP-AMPAR expression, we puffed AMPA onto the dendrites of RGCs. To confirm the identity of RGC cell type, we crossed Kcng4^cre/cre^ mice with a line in which a floxed STOP cassette prevents transcription of Td-tomato (Ai9, Jackson labs). This line labels α RGCs, both On and Off types (Duan et al., 2015; Krieger et al., 2017), in addition to at least one population of bipolar cell (Duan et al., 2014). We distinguished transient from sustained α Off RGCs by the presence of T type current as before. Off cells could be distinguished from On cells by the positioning of their dendrites, visualized by inclusion of Alexa 488 to the pipet solution. In addition, α On cells had faster and larger responses to AMPA than Off cells, presumably due to the proximity of their dendrites to the surface of the retina and the puff pipet. An exemplar IV relation of the AMPA puff response from an α Off transient cell is shown in figure 2B. Inspection of the raw currents shows that, as for light evoked currents, responses to AMPA did not rectify. The averaged currents generated by light driven synaptic responses, or direct activation of receptors with AMPA, are plotted at 8 different holding potentials in figure 2C. The responses are essentially linear, as indicated by a rectification index near 1.0 (Fig 2C, inset; Light: 0.90±.06, n=7; AMPA puffs: RI=1.00±.07, n=7).

We obtained similar results when measuring the IV relations of both light and AMPA evoked currents in α sustained Off RGCs (Fig. 2D,E). The amplitude of both types of responses were linear between holding potentials of −70 to +70 mV. The rectification index measured using either light or puffs of AMPA, were not significantly different, and both approached unity (Fig. 2F; RI_puff_: .87±.03, n=8; RI_light_: .94±.01, n=5, p=0.22). Taken together, it appears that endogenous and exogenous activation target the same, or largely overlapping AMPA receptor populations. In subsequent figures, data from both approaches are pooled.

### OHT selectively causes remodeling of AMPARs in α transient Off RGCs

We wanted to determine if chronic ocular hypertension (OHT) contributes to a change in AMPAR expression in α RGCs. We first focused on α Off type cells, using light and AMPA in combination with intracellular spermine to probe for changes. Mice were injected with microspheres in one eye (Fig 3A), while the other eye was either sham injected, or uninjected, serving as a control. Intraocular pressure (IOP) of both eyes was subsequently monitored (see methods for details). A clear elevation in IOP was observed in the bead-injected eye (~25% increase vs. control eye, p<0.0001), and persisted at day 18, the last time point that measurements were taken (Fig. 3B). Experiments were carried out on eyes following 14-21 days of OHT. An example of responses to light (Fig. 3C) and AMPA (Fig. 3D) in 2 α Off transient RGCs taken from the retina of a hypertensive eye are shown at 3 holding potentials. Note that the response to both AMPA and light are significantly reduced at positive holding potentials. The full IV relations of AMPA current evoked by light or AMPA puffs for α Off transient RGCs from both normal and OHT eyes are plotted in figure 3E. The rectification index for cells from normal eyes was 0.97±.04 (n=14) compared to 0.52±.03 in retinas from HT eyes (n=13), a highly significant difference (p<0.0001). Thus, α Off RGCs undergo a dramatic reorganization of synaptic AMPA receptors in response to 2-3 weeks of OHT.

**Figure 3.**
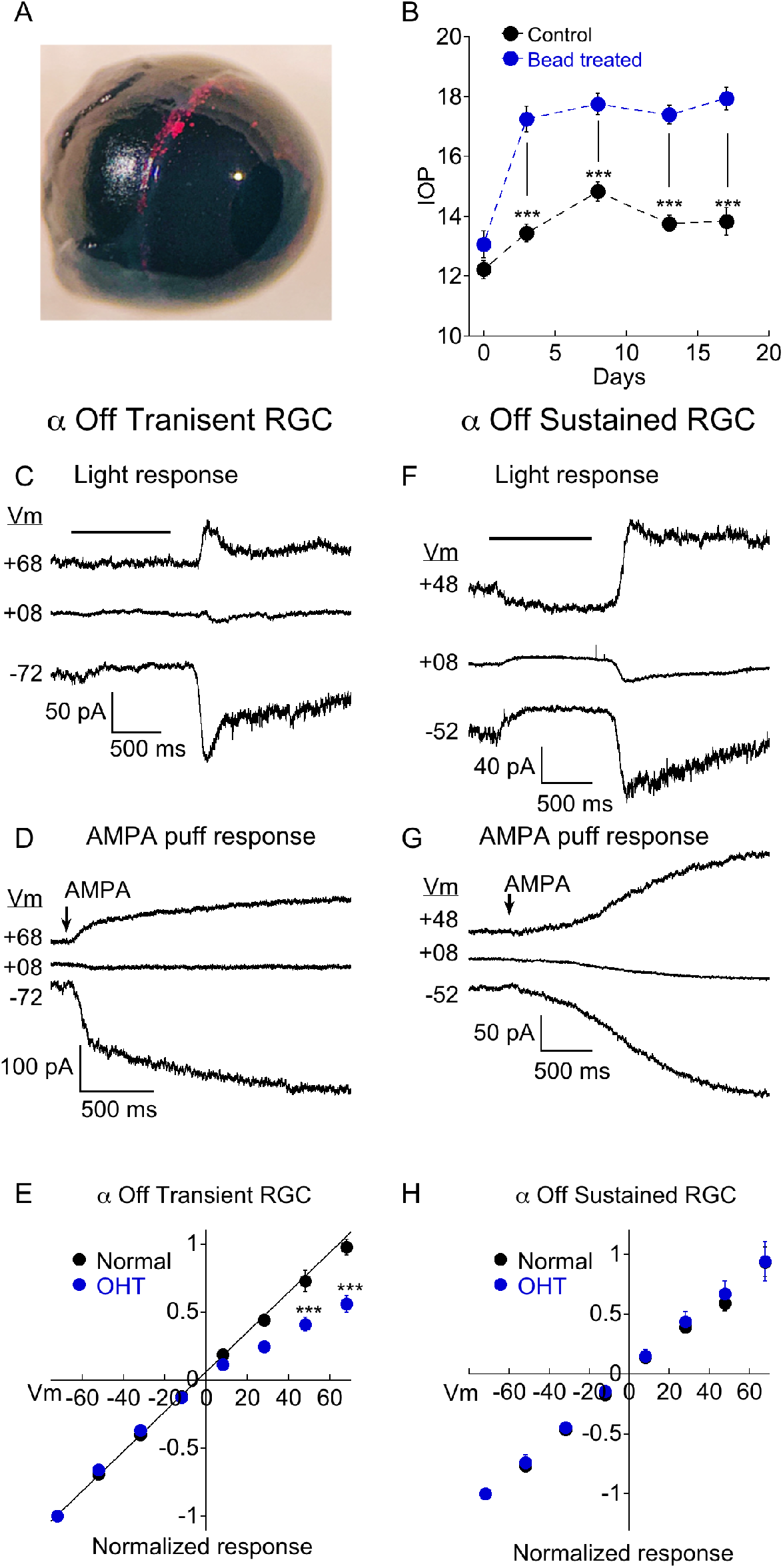
OHT drives remodeling of AMPARs in α Off transient RGCs, but not α Off sustained RGCs. A. Image confirming the presence of microbeads in the ciliary margin two weeks after microinjection of the eye. B. Measurements of intraocular pressure (IOP) of bead injected and sham injected eyes (control). Data are binned as follows: Day 0, 1-2 days before bead injection; day 3, 2-5 days after bead injection; day 8, 7-10 days; day 13, 12-15 days; day 17, 16-19 days. C. Light response of an α Off transient RGC at three holding potentials. The amplitude of the response at light Off is reduced at the positive voltage due to voltage dependent block of the CP-AMPAR component of the light response by spermine. D. Response of a different α Off transient RGC to AMPA at three holding potentials. The AMPA response is similarly reduced at positive voltage. E. IV relations for α Off transient RGCs from retinas with normal pressure and with OHT. IV relations obtained using AMPA and light have been pooled for both control (data from Fig 2; n=14) and OHT retinas (n=13). The difference was highly significant (*** indicates p<0.0001). F. Light response of an α Off sustained RGC from a hypertensive eye at three holding potentials. Responses at positive and negative holding potentials are equal in amplitude, despite the presence of spermine in the pipet solution, indicating a lack of CP-AMPAR expression. G. Responses to application of AMPA at positive and negative holding potentials are also of equal amplitude. H. Pooled IV relations comparing α Off sustained RGCs from retinas with normal IOP (data from Fig 2; n=13) and OHT retinas (n=11).

We performed similar experiments on α Off sustained cells. Recordings from sustained cells were obtained side by side with transient cells, from the same retinas. In contrast to transient cells, α Off sustained RGCs showed no evidence for increased expression of CP-AMPARs; responses to both light and AMPA puffs were nearly identical in magnitude at negative and positive holding potentials (Fig. 3F,G). The corresponding I-V relation for pooled α Off sustained RGCs was linear in retinas from mice with OHT, just as in retinas with normal pressure (Fig. 3H; normal: RI= .91±.06, n=13; OHT: RI=.83±.05, n=10; p=0.39). Thus both types of α Off cells express low levels of CP-AMPARs in normal retina, but transient Off RGCs selectively increase CP-AMPAR expression in response to elevated pressure.

### OHT also increases CP-AMPAR expression in α On RGCs

To probe for changes in AMPAR expression in α On RGCs driven by OHT, we activated AMPA receptors using two different strategies. As with α Off RGCs, we puffed AMPA directly onto the dendrites of α On RGCs. To activate receptors using endogenous synaptic input we made use of an optogenetic approach, crossing Cck-ires-cre mice to the Ai32 mouse line, which harbors ChR2 downstream from a floxed STOP cassette (Tien et al., 2017). This strategy confines expression of ChR2 to a single type of bipolar cell, cone bipolar type 6, which provides the majority of input to α On RGCs (Schwartz et al., 2012). To block the potential contribution of photoreceptor input to the optogenetic stimulation of type 6 bipolar cells, the mGluR6 agonist L-AP4 (10 μM) was included in the bath along with D-APV, picrotoxin and strychnine. Responses of two α On RGCs to either puffs of AMPA (Fig. 4A, left) or 1 ms optogenetic stimulation of type 6 bipolar cells (Fig. 4A, right), taken from eyes with normal IOP, are shown at three different holding potentials. Regardless of whether total AMPARs (using puffs of exogenous AMPA) or synaptic AMPARs (optogenetic stimulation) were examined, responses at positive voltages were nearly equal in magnitude to those recorded at negative voltages, indicating that CI-AMPARs were highly expressed at synaptic and extrasynaptic sites between type 6 bipolar cells and α On RGCs under normal conditions (Fig. 4C RI=.70±0.03, n=24).

**Figure 4.**
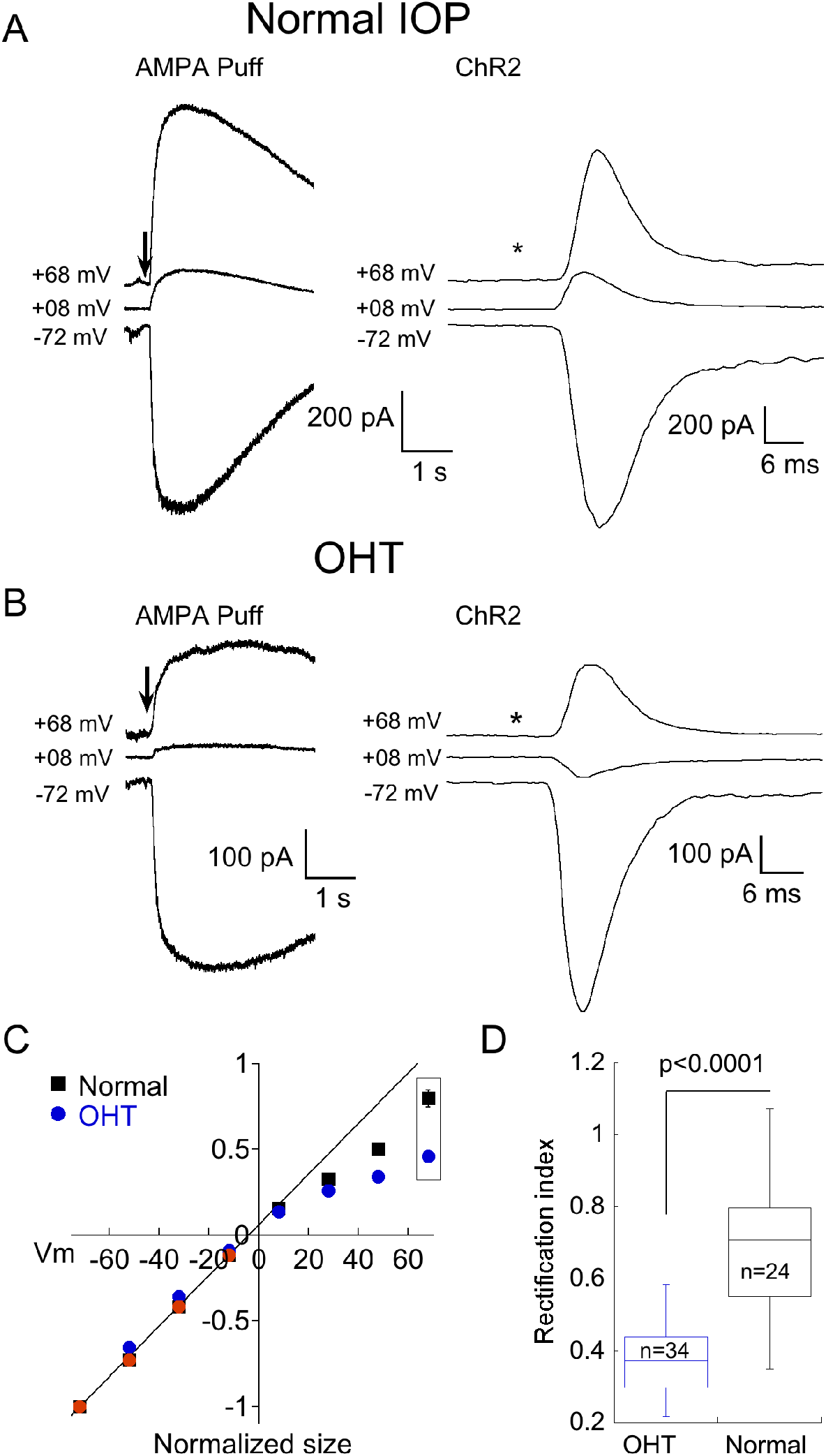
OHT also drives increased expression of CP-AMPARs in α On RGCs. A. Left, responses to 100 ms application of AMPA in an α On RGC from a retina with normal IOP at three holding potentials. Right, family of responses of an α On RGC to 1 ms stimulation of ChR2 expressed by type 6 bipolar cells (CCK-ires-cre:Ai32). B. As in (A), except that α On RGCs were from retinas with OHT. Note rectification of both AMPA-evoked current and the EPSC. C. Summary IV relations for α On RGCs from retinas with normal and OHT. Data are pooled from ChR2 and puff experiments. D. Plot summarizing rectification measured from the IV relations at +68 mV in (C) (boxed points) in OHT and normal retinas.

We next determined whether OHT induced remodeling of AMPARs in α On RGCs. Following 2-3 weeks of elevated IOP, CP-AMPAR expression was evaluated by measuring I-V relations with spermine present in the pipet solution as before. At positive holding potentials, responses to AMPA (Fig. 4B, left) and EPSC amplitude (Fig. 4B, right) were reduced, indicating a significant upregulation of CP-AMPARs. Increased expression of CP-AMPAR can be seen clearly in the IV relations comparing α On RGCs from normal and hypertensive eyes (Fig. 4C). The change in RI was highly significant, indicating robust insertion of CP-AMPAR (Fig. 4D, OHT; RI=.43±0.02, n=34, p<0.0001 vs. control eyes). Thus, AMPA receptor expression of both α On and α Off transient RGCs, but not α Off sustained RGCs, is altered by OHT.

### Editing of GluA2 plays a role in OHT-dependent remodeling of AMPARs

We next investigated the mechanism by which OHT increased CP-AMPAR expression. One potential mechanism is the replacement of AMPA receptors that contain the GluA2 subunit with receptors that do not. The second is a reduction in RNA editing of the Q/R site in the pore forming region of the GluA2 subunit, which would in turn reduce editing of glutamine to the positively charged arginine that prevents Ca^2+^ from passing through the channel. To distinguish between these two possibilities, we examined the effect of hypertension in a double mutant mouse line in which the enzyme responsible for RNA editing of GluA2, ADAR2, has been knockout out (MMRRC; Adarb1:Gria2, here abbreviated GluA2^R/R^). Since this is ordinarily a lethal mutation, a second mutation is the transgenic replacement of glutamine with arginine at the Q/R site, ensuring that AMPARs containing the GluA2 subunit cannot be Ca^2+^ permeable even in the absence of editing by ADAR2 (Higuchi et al., 2000). The responses of two α Off transient RGCs to light (left) or AMPA puffs (right) from the double mutant mouse line following 2-3 weeks of OHT are shown in figure 5A. Inspection of the raw data at positive and negative voltages shows a lack of inward rectification in either cell. Pooled I-V plots for α Off Transient RGCs from ocular hypertensive GluA2^R/R^ mice are shown along with I-V plots from OHT mice with normal GluA2 function (from Fig. 2) in figure 5C. Rectification of the IV for α transient Off RGCs in GluA2^R/R^ mice was nearly absent, and significantly different from bead treated mice with wildtype GluA2 (Fig. 5E, p<0.0001 vs. OHT GluA2 wt), but not significantly different from mice with normal IOP (p=0.265).

**Figure 5.**
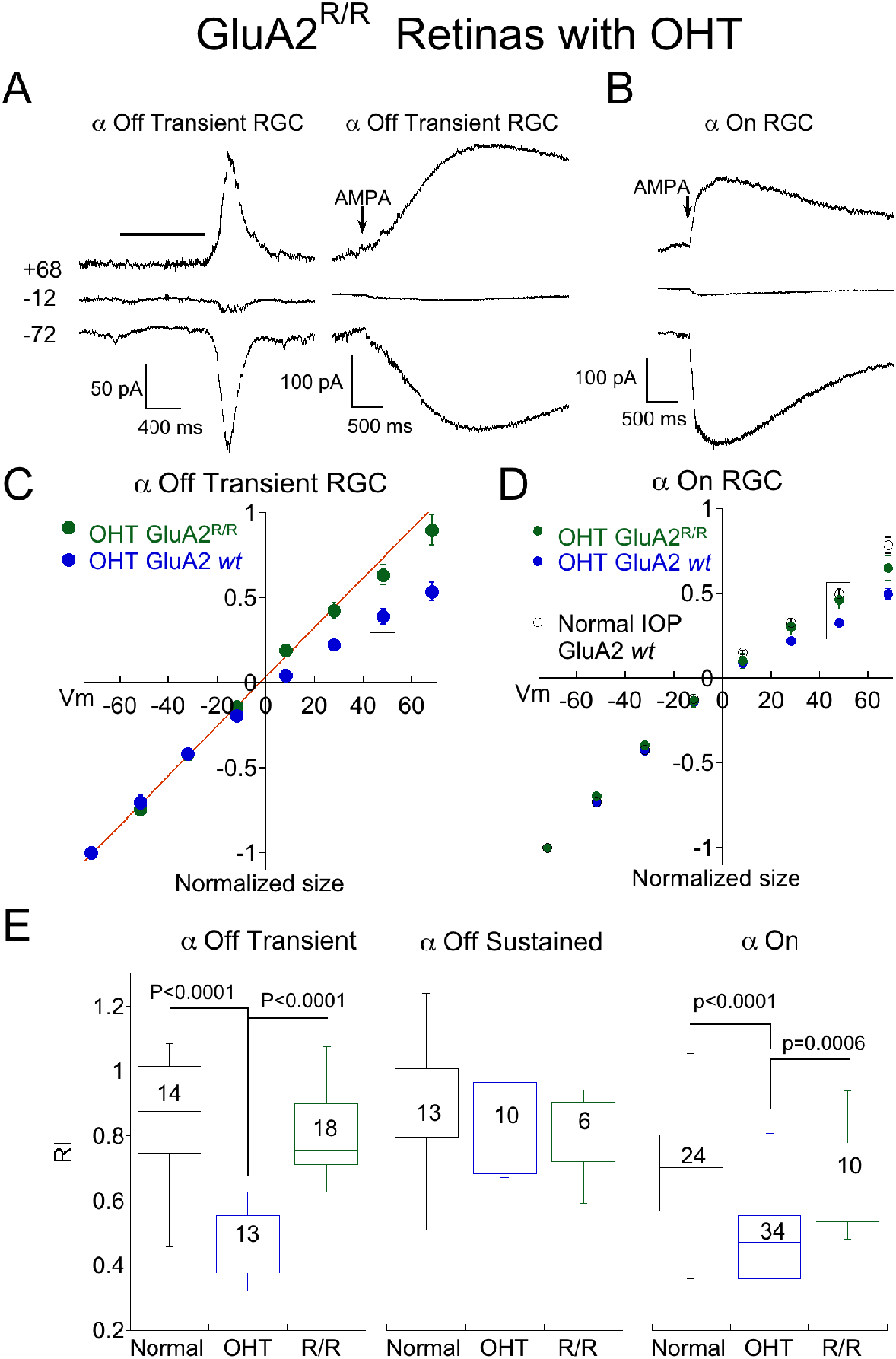
Genomic editing of the GluA2 Q/R site prevents remodeling of α On and transient Off RGCs by OHT. A. Recordings from two α Off transient RGC from the ADAR2^-/-^:GluA2^R/R^ mouse line, one showing responses to light (left) and the other showing responses to puffs of AMPA (right) at three holding potentials. Retinas were from OHT eyes. B. Responses to AMPA in an α On RGC from the same mouse line, also with OHT. C. Mean IV relationship of α Off transient RGCs for OHT retinas from wildtype mice (data replotted from Fig. 3) and ADAR2^-/-^:GluA2^R/R^ mice. The IV relations of α transient Off RGCs from mice with genomic editing of the Q/R site were linear, indicating a lack of CP-AMPAR expression. D. Mean IV relationship of α Off transient RGCs for OHT retinas from wildtype mice (data replotted from Fig. 4) and ADAR2^-/-^:GluA2^R/R^ mice. Also shown is the mean IV relation for wildtype mice with normal IOP. Genomic editing of the Q/R site linearized the IV relation of α On RGCs with OHT to nearly the same degree as cells from retinas with normal IOP. E. Summary rectification indices for all three α RGC types from normal and OHT retinas, and from ADAR2^-/-^:GluA2^R/R^ mice with OHT. In both α transient Off and α On RGCs, the effect of OHT on CP-AMPAR expression was reversed or reduced in the ADAR2^-/-^:GluA2^R/R^ mouse. For α sustained Off RGCs there was no significant difference between any condition.

Genomic editing of GluA2 also reduced rectification in α On RGCs of OHT mice, but the effect was more subtle. AMPA-evoked responses were smaller at positive voltages than at negative voltages, indicating the presence of rectification (Fig 6B). However, the I-V relation of α On RGCs of GluA2^R/R^ mice with OHT rectified substantially less than the I-V relation of α On RGCs from wildtype mice with OHT (Fig. 5D, wildtype I-V replotted from figure 4), and the difference in rectification between these two groups was highly significant (Fig. 5E; GluA2^R/R^ RI=.64±.04, n=10; wildtype RI=.47±.02, n=24; p=0.0006). Furthermore, there was no significant difference in IV rectification between α On RGCs from GluA2^R/R^ mice subjected to OHT and α On RGCs from wildtype GluA2 mice with normal eye pressure (Fig. 5E; p=0.84). Thus, CP-AMPAR expression induced by OHT can be largely eliminated by preventing a decrease in GluA2 editing in α On RGCs. However, there remains an additional component of CP-AMPAR expression in α On RGCs which is resistant to editing of GluA2, and is also present in retinas with normal IOP. It seems likely that this component of CP-AMPAR expression is contributed by GluA2 lacking AMPARs at type 6 BC-α On RGC synapses, and is present in α On but not α Off RGCs.

**Figure 6.**
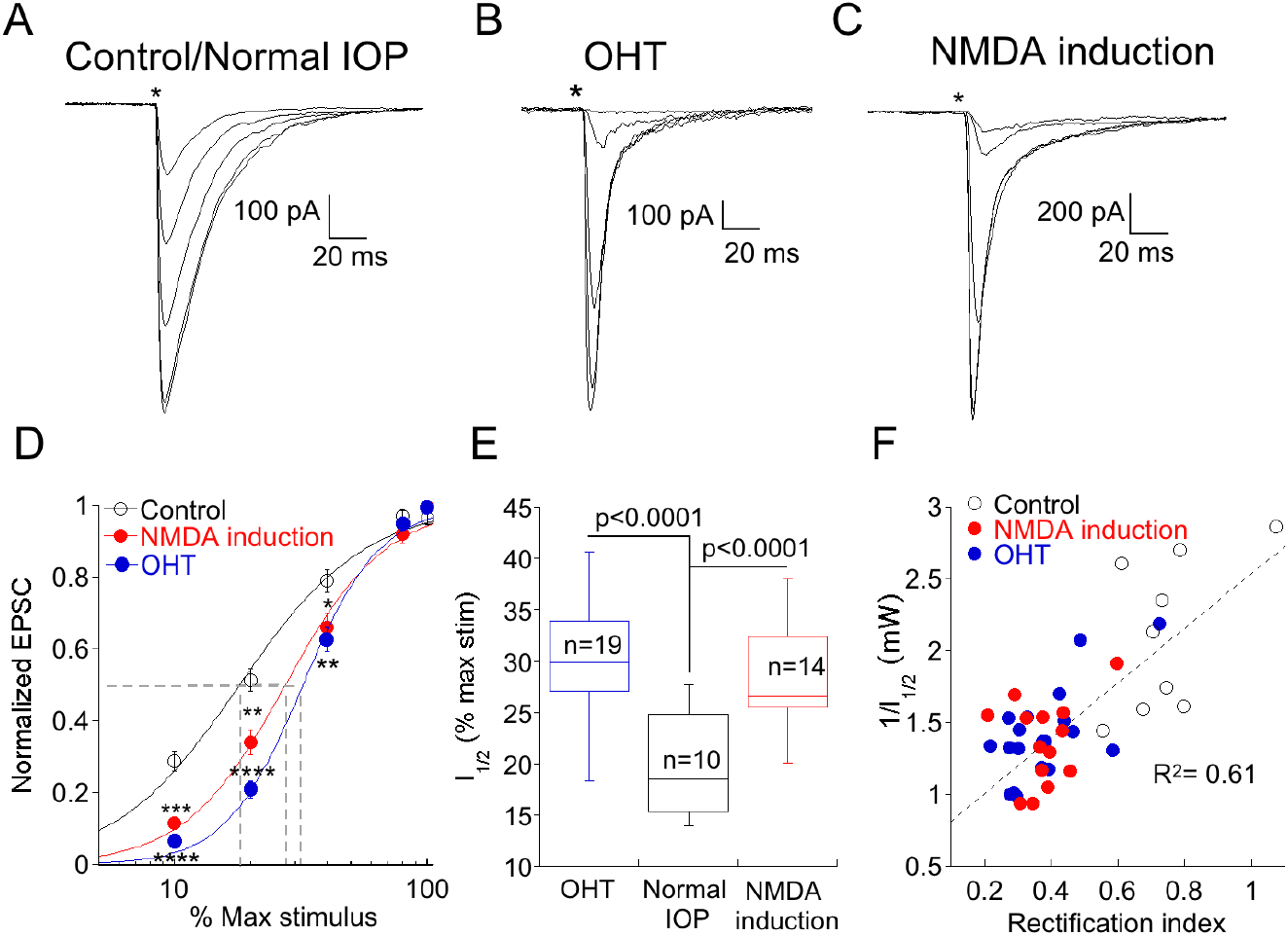
An increase in CP-AMPAR expression is associated with a decrease in synaptic gain at low stimulus intensities. A. Family of responses to a series of stimulus intensities in an α On RGC from a retina with normal IOP. The NMDA antagonist D-APV was present throughout the experiment to prevent NMDAR-dependent CP-AMPAR plasticity. B. Response of an α On RGC from an OHT retina to the same series of stimulus intensities. Responses are typical for α On RGCs from this condition as the response to weak stimuli are reduced or absent, compared with cells from retinas with normal IOP. C. Family of responses of an α On RGC from a retina with normal IOP, except that D-APV was present only during response measurements but not at other times. Under these conditions, RGCs showed a loss of sensitivity at low stimulus. D. Pooled data showing the normalized EPSC at each stimulus intensity under all three conditions. Fits are to the Hill equation with the following parameters: Control, slope=1.7, halfmax intensity=18% of maximum stimulus; NMDA induction, slope=2.2, halfmax intensity= 27% of maximum stimulus; OHT, slope=2.8, halfmax intensity=32% of maximum stimulus. Asterisks indicate significance for NMDA induction (upper asterisks) and OHT (lower asterisks) compared to control and are as follows: **** p<0.0001, *** p=0.0001, ** p=0.006, * p=0.04. 100% stimulus strength was 4.01 mW. E. Mean normalized response of α On RGC to a series of ChR2 stimulus strengths from normal and OHT retinas. E. Box plot of the ChR2 stimulus strength required to produce a half-maximal response in cells from normal eyes, normal eyes following induction of NMDAR plasticity, and hypertensive eyes. F. Correlation of synaptic gain, measured as the reciprocal of the half maximum intensity stimulus intensity, and CP-AMPAR expression, measured as the rectification index. Cells from all three conditions are indicated by separate symbols, but are treated as a single population for calculation of the linear regression.

As expected, no differences in CP-AMPAR expression were observed in α Off sustained RGCs from GluA2^R/R^ mice with elevated IOP compared with control mice with normal or elevated IOP.

### Elevation of CP-AMPAR expression is associated with a decrease in gain at the type 6 bipolar-α RGC synapse

A decrease in the gain of light responses of α On RGCs has been reported following elevation of IOP (Della Santina et al., 2013; Pang et al., 2015), but the mechanism or specific site responsible for the decrease has not been determined. Optogenetic stimulation of presynaptic type 6 bipolar cells affords the opportunity to isolate changes in the bipolar cell-ganglion cell synapse that are driven by OHT. We stimulated type 6 bipolar cells in normal and OHT eyes, varying the voltage of the input driving the LED to control the light intensity and subsequent bipolar cell type 6 depolarization, and recorded the resulting EPSCs in α On RGCs to generate an “intensity”-response function (Fig. 6A). When the same light intensities were used to evoke EPSCs in α On RGCs from hypertensive eyes, responses were reduced or absent at lower stimulus intensities compared to normal retinas (Fig. 6B). However, responses between the two groups were not significantly different at high stimulus intensities. The relationship between the LED input voltage and α On EPSC was highly repeatable across cells from normal and OHT eyes, and a good fit to the response relationship for both conditions could be obtained using a standard Hill function (Fig. 5D, control vs. OHT). Thus, OHT decreases synaptic gain at lower stimulus strengths, and increases CP-AMPAR expression at the type 6 bipolar-α On RGC synapse, raising the possibility that the two effects might be causally related.

To gain more insight into this possibility, we sought an alternative model in which CP-AMPAR expression is increased in the absence of OHT. High frequency stimulation of the bipolar-α On RGC synapse causes rapid insertion of CP-AMPARs through a mechanism that involves NMDARs (Jones et al., 2012). In the present study, experiments were carried out in room light, and D-APV was always present in the bath to block insertion of CP-AMPARs. In the absence of the NMDAR antagonist, the rectification index was 0.45±.03 (n=35, p<0.0001 compared to retinas treated with D-APV), indicating that room light alone is sufficient to cause substantial insertion of CP-AMPARs when NMDARs are not blocked. We therefore asked whether synaptic gain was decreased at synapses where NMDAR-dependent plasticity had been induced. Similar to synapses from retinas with OHT, we observed a decrease in EPSC amplitude at low stimulus intensities (Fig. 6C). The relationship between EPSC amplitude and stimulus intensity was similar when CP-AMPAR expression was elevated either by NMDAR-dependent plasticity or by OHT (Fig. 6D). Both OHT and NMDAR plasticity raised the stimulus intensity required to evoke 50% of the maximum EPSC (I_1/2_) compared to cells from retinas with normal IOP in the presence of D-APV (Fig. 6E). To further quantify the relationship between CP-AMPAR expression and synaptic gain, we plotted the rectification index of each cell in all three groups vs. the reciprocal of I_1/2_ (Fig. 6F). Surprisingly, the relationship between these two parameters was highly correlated across groups (R^2^ = 0.61, p=1.7X10^-8^); α On RGCs with high levels of CP-AMPAR expression had lower synaptic gain, and this relationship was agnostic to the conditions the cell had experienced. These experiments support the idea that a decrease in light sensitivity associated with elevated ocular pressure is a consequence of CP-AMPAR expression at the type 6 bipolar-α On RGC synapse.

### Postsynaptic manipulations of α On RGCs can alter transmitter release from type 6 bipolar cells

What can account for the decrease in synaptic gain associated with elevated expression of CP-AMPARs? Changes in synaptic strength can have a presynaptic or postsynaptic locus, or both. As a starting point, we measured paired pulse ratios (PPR) a standard method for assessing changes in release probability. When using stimulus strengths that produced maximum EPSC amplitudes, α On RGCs from retinas with normal IOP and low CP-AMPAR expression (Fig. 7A, left) exhibited significant paired pulse depression (PPD) at short intervals (50 ms), suggesting that, transmitter release probability from the type 6 BC is high in response to strong ChR2 stimulation (Fig. 7A, right). At longer intervals, depression decreased, presumably due to replenishment of released vesicles. The rate of recovery could be adequately fit by the sum of a fast (τ_fast_=94 ms) and slow (τ_slow_=2.8 s) exponential, similar to the kinetics of recovery from depletion at rod bipolar cell terminals (Singer and Diamond, 2006; Wan and Heidelberger, 2011), and at climbing fiber presynaptic terminals in cerebellum (Dittman and Regehr, 1998).

**Figure 7.**
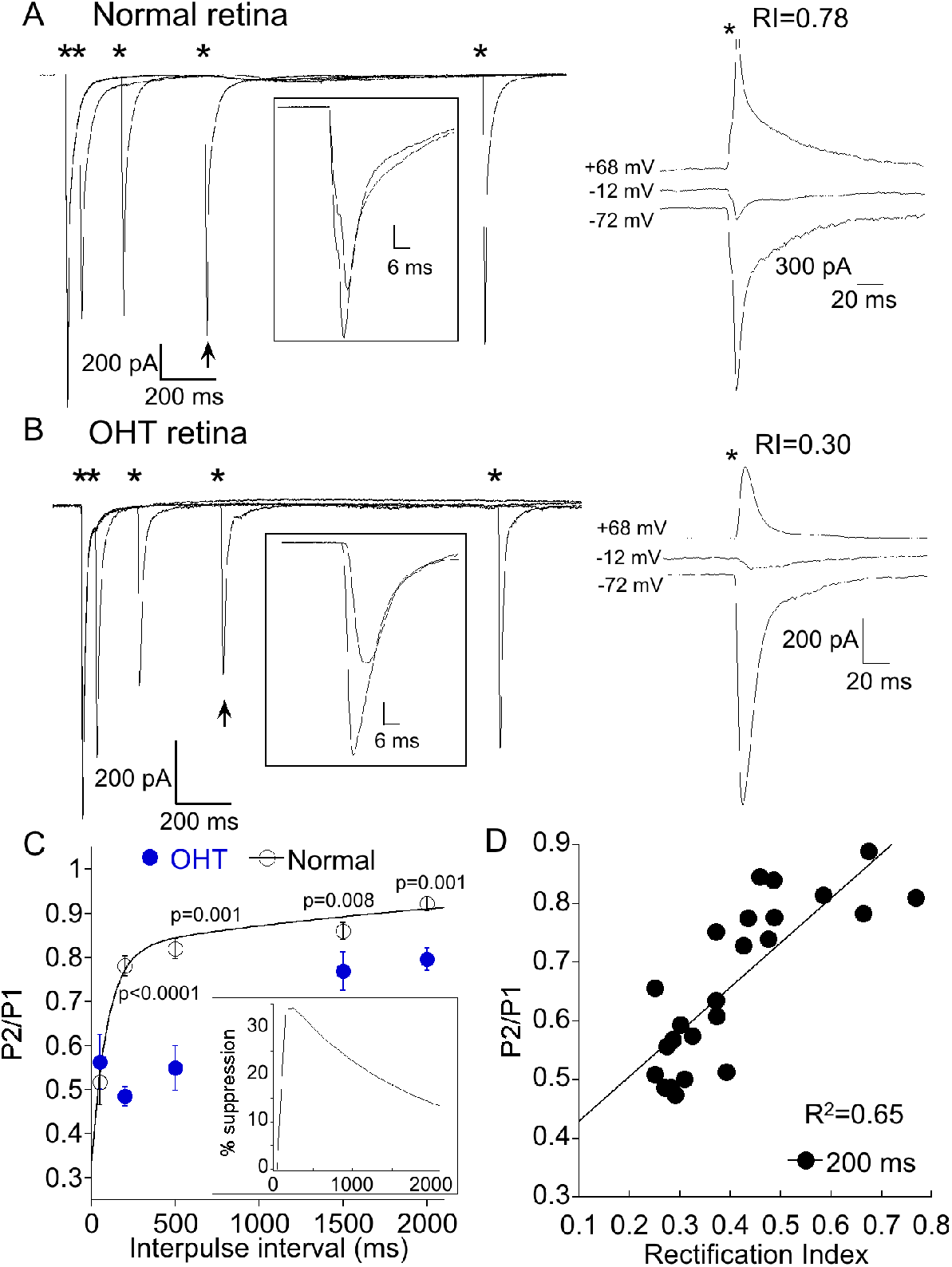
AMPAR remodeling is associated with delayed short-term synaptic depression. A. Left, responses of an α On RGCs from a retina with normal IOP to paired 1 ms, 8.0 mW ChR2 stimulation at intervals indicated by the asterisks. Inset: Overlay of the first response with the response obtained 500 ms later (indicated by arrow). Recovery of the EPSC is nearly complete after 500 ms. Right, IV relation of the same cell shows minimal amount of rectification, indicating low CP-AMPAR expression. B. As in (A), but the α On RGC was from an OHT retina. Rectification at positive voltage is pronounced in this cell. Inset shows reduced recovery of the EPSC at this time point. C. Summary of paired pulse depression at the indicated pulse intervals of α RGCs from OHT (n=17) and normal retinas (n=14). The sum of two exponentials, A_fast_*exp(-t/τ_fast_) and A_slow_*exp(-t/τ_slow_), was used to fit the recovery of the EPSC for α RGCs from retinas with normal IOP. The fits were extrapolated to 15 seconds following the first stimulus, which represented full recovery of the EPSC. Inset: Time course of the delayed suppression of the EPSC obtained from the difference of the mean fits to the normal and OHT conditions. D. Correlation of rectification index with paired pulse ratio at 200 ms. The delayed paired pulse ratio was highly correlated with CP-AMPAR expression, with delayed suppression of the EPSC (i.e., lower P2/P1 ratio) greater for cells expressing higher a higher fraction of CP-AMPARs (low RI) at synapses (p=2.0X10^-6^).

We performed paired pulse experiments on α On RGCs from OHT eyes with elevated CP-AMPAR expression (Fig. 7B). At high stimulus intensities, paired pulse depression at an interval of 50 ms was similar for normal and OHT eyes. At this interval the ratio of the second to first EPSC was 0.53±.06 for normal eyes, and 0.51±.05 for OHT eyes (p=0.83), suggesting that the initial release probability at type 6 bipolar cells under both experimental conditions is essentially the same. When we examined the PPR at longer intervals, we obtained a surprising result: Subsequent EPSCs were dramatically reduced compared with EPSCs at similar time intervals in normal retinas (compare responses indicated by arrows and insets). In normal retinas, the EPSC evoked at 200 ms recovered to 78±2% of the first EPSC, but only to 49±3% of the first EPSC in OHT eyes (Fig. 7C, p<0.0001). The slowed time course of the recovery from paired pulse depression of α On RGCs in hypertensive eyes can be explained by the addition of a delayed component of EPSC suppression superimposed on the recovery of EPSC from vesicle depletion. The delay in EPSC suppression varied slightly from cell to cell, but reached a peak at 150-250 after the initial stimulus (Fig. 7C, inset). At an interval of 2 seconds, the EPSC in OHT eyes was still significantly suppressed relative to control animals. This delayed suppression of the EPSC persisted in the presence of the GABA_c_ antagonist TPMPA, ruling out inhibitory feedback from amacrine cells as a potential mechanism.

To quantify the relationship between this form of delayed synaptic depression and CP-AMPAR expression, we compared the paired pulse ratio (PPR) at 200 ms with the RI for each cell. We pooled cells from normal and OHT eyes, using only RI as a variable. The RI was strongly correlated with the PPR at 200 ms (Fig 7D, R^2^=0.65; p=2.0X10^-6^). Below we consider the possibility that the first stimulus and EPSC provide a trigger that leads to a subsequent rapid and brief suppression of excitatory transmission, perhaps via a retrograde signaling pathway. CP-AMPARs may play a role in this process by providing a route for Ca^2+^ entry intro the postsynaptic cell to initiate the signaling pathway.

If Ca^2+^ plays a role in the initiation of a postsynaptic signal that is retrogradely communicated to type 6 bipolar cell terminals, then inclusion of BAPTA in the pipet solution to rapidly buffer Ca^2+^ might reduce or eliminate this signal as has been shown elsewhere for retrograde signaling. In support of this idea, dialyzing 20 mM BAPTA into α On RGCs with high CP-AMPAR expression (RI< 0.4) prevented delayed synaptic depression (Fig. 8A). The kinetics of recovery in these cells (Fig. 8C, τ_fast_=81 ms, τ_slow_=2.0 s) were similar to α On RGCs that had low CP-AMPAR expression (Fig. 7). Retrograde signaling is often mediated by release of endocannabinoids, which act on presynaptic CB1 receptors. To determine if CB1 receptors play a role in the delayed suppression of EPSCs observed at α On RGCs, we applied 2 μM AM251, a CB1 antagonist, to the retina and measured the paired pulse ratio. To ensure that AM251 had sufficient time to penetrate the tissue, and because washing out of AM251 is slow, the antagonist was continuously perfused into the retina, and cells with high CP-AMPAR expression were chosen. Incubation of retinas with AM-251 also prevented delayed EPSC suppression (Fig. 8B). The kinetics of recovery from paired pulse depression were similar to cells recorded with BAPTA or cells with low CP-AMPAR expression (τ_fast_=58 ms, τ_slow_=2.1 s). The difference in PPD at 200 ms between high CP-AMPAR expressing cells recorded with EGTA and those recorded with BAPTA in the pipet solution or with AM-251 in the bathing solution was significant (Fig. 8D). Thus either strong buffering of Ca^2+^ or block of CB1 receptors eliminated the delayed suppression of EPSCs, leaving behind an exponential recovery from transmitter depletion that was similar to the recovery observed in α On RGCs with low expression of CP-AMPARs.

**Fig 8.**
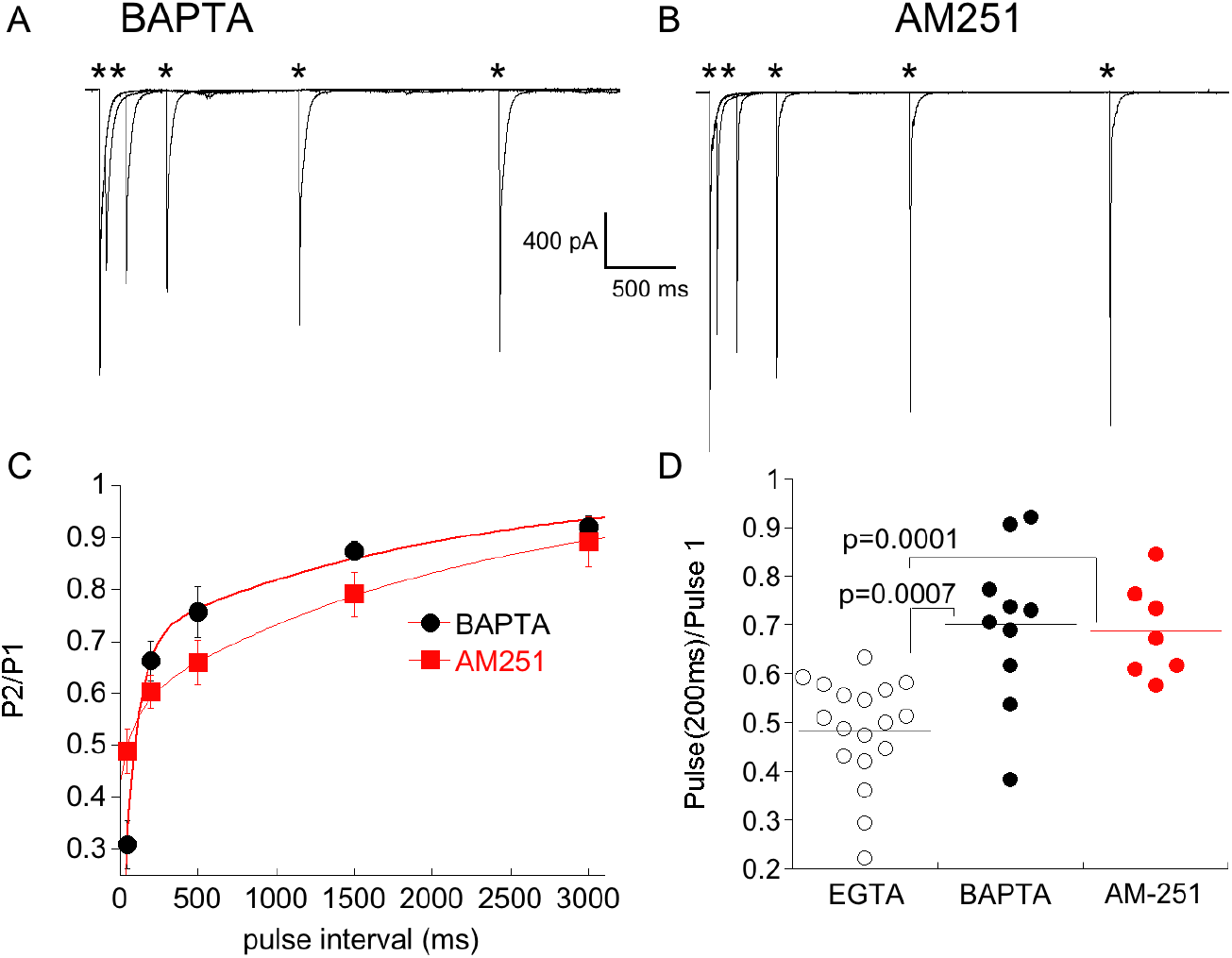
Postsynaptic buffering of Ca^2+^ or blocking CB1 receptors abolishes synaptic depression at CBC6→α On RGC synapse. A. Responses from an α On RGCs to paired, 1 ms stimuli at the indicated stimulus intervals. The internal solution contained 20 mM BAPTA. B. Responses of an α On RGCs to paired, 1 ms stimuli as in (A) with 2 μM AM251 added to the bathing solution. Summary plot of the paired pulse ratio under the conditions shown in (A) and (B). Cells recorded with the internal solution containing BAPTA and AM251 are from retina with normal IOP, but were induced to undergo NMDA-dependent plasticity by omitting D-APV. Only α On RGCs that displayed significant rectification (RI<0.4) were included for analysis. C. Time course of recovery from paired pulse depression in α On RGCs recorded with BAPTA in the pipet solution, or with EGTA in the pipet and AM251 in the bath. Paired pulse depression decreased monotonically, with no evidence of a delayed increase in depression despite signifcant expression of CP-AMPARs in these cells. D. Statistical comparison of paired pulse ratio at an interval of 200 ms in cells recorded with EGTA (group of α On RGCs from Fig. 7C), BAPTA, or with AM-251 in the bath.

## Discussion

In this study, we show that elevation of IOP via injection of microbeads into the anterior chamber, a widely used experimental model for glaucoma (Sappington et al., 2010; El-Danaf and Huberman, 2015; Pang et al., 2015; Risner et al., 2018), leads to upregulation of CP-AMPARs in RGCs. This form of AMPAR remodeling is cell-type specific, completely absent in α Off sustained RGCs, robust in α Off transient RGCs, and more modest in α On RGCs. Furthermore, we find that remodeling of AMPARs is absent in RGCs from mice in which the Q/R site is genetically altered to arginine. This finding supports the idea that a reduction in editing by ADAR2 at the Q/R site is responsible for the increase in CP-AMPAR expression associated with chronic OHT. Robust AMPAR remodeling of α transient Off RGCs observed in the present study is in line with the notion that this cell type is particularly susceptible to OHT (Della Santina and Ou, 2017). Within the first 7 days, α transient Off RGCs undergo extensive dendritic pruning and loss of synaptic contacts, judged by a reduction in PSD-95 puncta (Della Santina et al., 2013; El-Danaf and Huberman, 2015; Ou et al., 2016). Physiological changes in parameters such as stimulus-evoked spike frequency, response sensitivity receptive field size, and spontaneous activity have been documented for α transient Off RGCs (Della Santina et al., 2013; Pang et al., 2015; Ou et al., 2016) and they are observed prior to changes in structure. The finding that CP-AMPARs are upregulated during the same time period raises the possibility that Ca^2+^ influx through abnormally high levels of these receptors plays a role in degradation of dendrites.

### Is remodeling of AMPAR a consequence of low expression of ADAR2 or downregulation of GluA2?

Remodeling of AMPARs is a well established consequence of a wide spectrum of neuronal dysfunction, including drug dependence (Wolf, 2016), seizures (Rakhade et al., 2008; Lippman-Bell et al., 2016), and ischemia (Noh et al., 2005; Dias et al., 2013; Hwang et al., 2013). While the evidence for increased expression of CP-AMPAR is often clear, the underlying mechanism, either a decrease in ADAR2 levels and consequently of GluA2 editing, or a downregulation of GluA2 transcription, is not. Ultimately an understanding of the underlying mechanism has implications for treatment strategies. Downregulation of ADAR2 and subsequent increases in CP-AMPARs has been identified as a key step in the progression of motor neuron loss in ALS (Kwak and Kawahara, 2005; Hideyama et al., 2012; Lorenzini et al., 2018; Yamashita and Kwak, 2018). Furthermore, deletion of ADAR2 in motor neurons is sufficient to phenocopy ALS, and can be rescued by genomic editing of the Q/R site (Hideyama et al., 2010). To establish a causal role in glaucoma, it would be instructive to determine whether the structural and functional loses associated with OHT are rescued in the GluA2^R/R^ mouse line, where AMPAR remodeling is not observed.

OHT has also been shown to increase CP-AMPAR in RGCs of rat retina through a mechanism involving TNF-α (Cueva Vargas et al., 2015). However, TNF-α appears to increase CP-AMPAR expression by a down regulation of GluA2, rather than altering ADAR2 editing. In that study, the authors reduced outflow through the episcleral veins, resulting in an increase in IOP that was 2-fold higher than the IOPs achieved in the present study. Perhaps a larger increase in IOP recruits additional mechanisms of AMPAR remodeling. Furthermore, the authors of that study did not identify the affected RGC subtypes, leaving open the possibility that OHT activates different mechanisms of AMPAR remodeling, depending upon the specific RGC subtype.

In contrast to maladaptive CP-AMPAR expression under pathological conditions, adaptive changes in CP-AMPAR expression are an essential component of many forms of long and short-term plasticity that involve recruitment of existing or newly synthesized GluA2-lacking receptors (Ju et al., 2004; Clem and Barth, 2006; Sutton et al., 2006). In α On RGCs, a rapid NMDA-dependent increase in CP-AMPAR expression has been described previously (Jones et al., 2012) and was revealed in the present study by comparing CP-AMPAR expression in separate populations of α On RGCs either in the presence or absence of NMDAR blockers. Even under conditions of NMDAR block, α On RGCs expressed significant levels of CP-AMPARs, perhaps in response to lack of synaptic activity, a form of synaptic scaling described at a number of synapses (Isaac et al., 2007), including this one (Xia et al., 2007). This component of CP-AMPAR expression was spared in the GluA2^R/R^ mouse, implying that it does not result from a decrease in GluA2 editing. Thus, it appears that both known mechanisms of AMPAR remodeling converge onto α On RGCs, one as a result of pathological OHT, the other due to a physiologically relevant stimulus.

### A decrease in synaptic gain is associated with elevated CP-AMPAR expression

The observation that both NMDA-dependent plasticity and OHT reduced synaptic gain in α On RGCs led us to investigate the possibility that CP-AMPAR expression and gain were linked. To drive synaptic input to α On RGCs, we used an optogenetic approach. This eliminates confounding effects of OHT on upstream circuit elements. It also provides higher temporal resolution than other approaches, The ChR2-EGFP fusion protein was expressed in type 6 bipolar cell, which provides the majority of input to α On RGCs (Schwartz et al., 2012; Tien et al., 2017), while α Off RGCs appear to receive input from different cohorts of bipolar cells, (Yu et al., 2018) allowing for only a minority of synaptic inputs to be stimulated optogenetically. We examined the relationship between the strength of the stimulus driving ChR2 and the EPSC (Najac and Raman, 2017), and found this relationship to be quite repeatable across cells. One reason for this might be the high degree of convergence of synaptic connections from CB6 input onto individual α On RGCs (Freed et al., 1992; Kerschensteiner et al., 2009; Morgan et al., 2011; Schwartz et al., 2012), allowing for the smoothing of variability in ChR2 expression between bipolar cells.

The delayed suppression of EPSCs that we observed in OHT and NMDA-treated retinas shares some properties of depolarization-induced suppression of inhibition (DSI) (Pitler and Alger, 1992, 1994) and depolarization-induced suppression of excitation (DSE) (Kreitzer and Regehr, 2001b). Both DSI and DSE transiently suppress synaptic transmission with a short latency following depolarization. Suppression is due to retrograde release of endogenous cannabinoids which bind to presynaptic CB1 cannabinoid receptors to reduce Ca^2+^ influx into synaptic terminals (Kreitzer and Regehr, 2001a; Wilson and Nicoll, 2001; Ohno-Shosaku et al., 2002; Chevaleyre and Castillo, 2004). Release is stimulated by a rise in postsynaptic Ca^2+^ through voltage gated channels. In the present study, depression was initiated by a single stimulation of transmitter release, rather than postsynaptic depolarization, raising the possibility that Ca^2+^ influx through open CP-AMPARs is sufficient to turn on production of a retrograde messenger such as 2-AG or anandamide, endogenous activators of CB1 receptors (Piomelli, 2003). Local increases in Ca^2+^ generated by influx through CP-AMPARs is sufficient to stimulate transmitter release from A17 amacrine cells in the absence of voltage gated Ca^2+^ channels (Chavez et al., 2006), and stimulation of CP-AMPARs at mossy fiber-CA3 interneuron synapses initiates a presynaptic form of LTD (Lei and McBain, 2004). CB1 receptors are expressed in the inner retina in a number of species (Straiker et al., 1999; Yazulla et al., 1999), and cannabinoids have been shown to modulate synaptic input to RGCs (Middleton and Protti, 2011) and decrease Ca^2+^ currents in presynaptic bipolar cells (Straiker et al., 1999).

## Methods

### Animals

Mice of either sex were used in this study. Mice were obtained from The Jackson Laboratory. For experiments in dark adapted retinas, C57Bl/6j were used. For identification of α RGCs we crossed the Kcng4^cre^ (029414) with a Td-Tomato Cre reporter line (Ai14). For channelrhodopsin2 (ChR2)-mediated depolarization of Type 6 bipolar cells, we crossed CCK^cre^ (012706) with a line that expressed ChR2 following cre-mediated excision of an upstream STOP sequence (Ai32). The ADARB1^-/-^ Gria2^R/R^ mouse line (Adarb1^tm1phs-^ Gria2^tm1.1phs^/Mmnc) was obtained from the MMRRC (034679-UNC). These mice will be referred to as GluA2^R/R^.

### Bead injection

All procedures were in accordance with the animal care guidelines for the University of Nebraska Medical Center Institutional and Animal Care Use Committee. Animals were anesthetized with isoflurane, pupils were dilated with 1% tropicamide ophthalmic solution (Bausch & Lomb), and anesthetic drops (0.5% proparacaine hydrochloride; Bausch & Lomb) were applied to one eye. The anterior chamber was injected with 10 μm polystyrene microbeads (cat #F8834, Invitrogen). The bead suspension was concentrated by centrifugation of 200 μl of solution followed by removal of 150 μl supernatant. For delivery of beads, glass tubing (type 7052, King Glass) was pulled to a diameter of 50 μM using a vertical puller (Narishige). The pipet was type filled with 1-2 μl of hyaluronate (Provisc, Alcon) followed by 1-2 μl of bead solution. Injection of hyaluronate before removing the pipet sealed the entry hole of the pipet and prevented efflux of the beads. Beads were ejected using a manual microsyringe pump (World Precision Instruments). Following the injection, the antibiotic ciprofloxacin was applied to the eye. Control retinas were either from uninjected or sham injected eyes. No statistical difference in results were observed between these two groups. IOP measurements were made in both eyes with the TonoLab tonometer beginning 1-2 days prior to bead injection and once every 3 days for 18 days after injection. IOP was measured in anesthetized animals.

### Preparation of the retina

For experiments that measured light responses, mice were dark adapted overnight before killing them with CO_2_ inhalation followed by cervical dislocation. Retinas were isolated under dim red light and retinas were incubated in a solution containing collagenase and hyaluronidase dissolved in oxygenated (95% O_2_ and 5% CO_2_) Ames media (Sigma-Aldrich) at room temperature 15-30 minutes to aid in penetration of the inner limiting membrane (Schmidt and Kofuji, 2011). Retinas were then mounted in the recording chamber and held in place with a slice anchor (Warner Instruments) and perfused with Ames bubbled with 95% O_2_ and 5% CO_2_ at a rate of 4-6 ml/min. For optogenetic and puffing experiments, mice were dark adapted for one hour prior to sacrifice, and all manipulations were carried out in room light.

### Imaging

10 μM Alexa488 Hydrazide (ThermoFisher; catalog #A10436) and 0.5% Neurobiotin (Vector Labs; catalog #SP-1120) were added to the normal internal solution. After recording, retinas were fixed in 4% paraformaldehyde for 30 minutes, washed in PBS 3 times, and then incubated in a blocking solution of 5% donkey serum, 1% triton X-100 and 0.5% DMSO in PBS for 1 hour at room temperature. Retinas were then incubated in blocking solution and goat anti-ChAT primary antibody (1:100; Millipore Sigma, catalog #AB144P) and Texas Red-conjugated streptavidin (Vector Labs; catalog #SA-5006) for 5 days at 4°C. Retinas were then rinsed 3X and incubated for 2 hours in Texas red-conjugated donkey anti rabbit secondary (1:100; ThermoFisher; catalog #PA1-28662) at room temperature, washed 3X in PBS, and mounted with Slowfade antifade (ThermoFisher catalog #S2828). Images were taken with a Nikon A1 confocal microscope with an oil immersion 63X objective. Image dimension was 512X512 pixels (pixel size: 0.41 μm). Z scan resolution was 1 μm. Images were compiled using Fiji (ImageJ) software. *Patch clamp recording*. The retina was viewed on a video monitor using infrared illumination and a CCD camera (COHU Electronics) mounted to an Olympus Bx51 microscope equipped with a water-immersion 40X objective. In dark adapted conditions, transient and sustained Off α RGCs were targeted based on their large (~20 μm) somas, response to light, and the presence or absence of a T type Ca^2+^ current. For experiments using puffs of AMPA, a Kcng4^cre^:Ai14 reporter line was used to identify α RGCs, and further classification into subtypes was carried out by filling cells to visualize the depth of dendrites (On vs. Off) and measurement of a T type current (Off transient vs. Off sustained). Pipettes (tip resistance 5–7 MΩ; 1.5 mm OD, WPI) were filled with a cesium gluconate solution containing the following (in mM): 123 Cs gluconate, 8 NaCl, 1 CaCl2, 10 EGTA, 10 HEPES, 10 glucose, 5 ATP, .01 Alexa 488 (puff and optogenetic experiments), and 100–500 μM spermine (pH 7.4; 290 mOsm). To isolate the AMPAR-mediated EPSC, strychnine (1 μM), picrotoxin (100 μM), and D-AP5 (50 μM) were added to Ames media. TPMPA (50 μM) was included to blocked inhibitory feedback onto bipolar cells mediated by GABA_C_ receptors. All chemicals were purchased from Sigma-Aldrich or Tocris Bioscience. Cells were voltage clamped at −60 mV. Current voltage relations were obtained by stepping cells from −60 mV to +80 mV in 20 mV increments. Cells were held at 0 mV between steps. Holding potentials were corrected for a 12 mV junction potential off line, and so voltages ranging from −72 to +68 mV are reported here. Series resistance, typically measuring 8–20 MΩ, was not compensated for. Recordings were discarded if series resistance was >20 MΩ. Recordings were obtained with an Axopatch 200B (Molecular Devices) using AxoGraph X acquisition software and digitized with an ITC-18 interface (Heka Instruments).

### Light activation of AMPA currents

The light source for generating responses in the dark adapted retina was a 20 W halogen lamp focused through a 40X objective. An interference filter (peak transmittance at 500 nm) and neutral density filters were inserted in the light path to control the intensity and wavelength of light stimulation, and a shutter (Uniblitz; Vincent Associates) was used to control the duration of the stimulation, typically 1 s. The intensity of the unattenuated light stimulus, measured using a ratiometric spectrophotometer (Thor Labs) was 4.9X10^5^ R*/rod·s at 490 nm, assuming a collecting area of 0.5 μM per rod (Field and Rieke, 2002).

### Optogenetic activation of AMPA currents

ChR2 was activated with a 1 ms activation of a 490 nm LED (Sutter Instruments). To measure responses over a range of type 6 bipolar cell membrane depolarizations, the voltage driving the led was regulated using a computer controlled analog input to the LED. We used 5 voltages (0.25 V, 0.5 V 1 V, 2 V, 2.5 V). The LED intensity at each voltage, measured using a spectrophotometer (Thor Labs), was stable over the time period over which these experiments were performed. To prevent photoreceptor inputs, L-AP4 (10 μM) was included in the bath along with the standard antagonist cocktail.

### Activation of AMPA currents with AMPA

To apply AMPA we used a second pipet coupled to a positive pressure device (Picospritzer, Parker). For α On RGCs, the puffer pipet was positioned at the surface of the retina. For α Off RGCs, the puffer was advanced into the retina. Even before identification of cell type was confirmed anatomically, α On cells could be readily identified by the more rapid onset and decay of AMPA responses compared with Off α RGCs, whose dendrites ramify more deeply in the IPL. The continuous flow of Ames media from a locally positioned capillary tube (ID 200 μm, Polymicro Technologies) dramatically increased the rate of diffusion of AMPA from the tissue following each puff. Puffs of AMPA were applied at 20 second intervals to insure sufficient time for washout from the tissue.

### Experimental design and statistical analysis

Analysis was performed using AxoGraph X and KaleidaGraph (Synergy Software) software. The rectification index (RI) we calculated by first fitting the currents obtained at negative holding potentials with a linear regression. The RI was the ratio of the amplitude of the current evoked at +48 mV was to the current amplitude at the same voltage predicted by the regression. We chose +48 mV rather than +68 mV as the test voltage, as some cells showed a reduced spermine block at the more positive voltage (Bowie and Mayer, 1995). Statistical significance was determined using the Wilcoxon-Mann-Whitney test. Error bars indicate the SEM. Details regarding the number of cells for each condition can be found in the figure legends.

## Acknowledgements

We thank Dr. Wallace Thoreson for valuable discussions and critical reading of the manuscript, and Cody Barta for technical assistance.

